# Single-strain mobilome sequencing quantifies bacterial genetic response to stress, including activity of IS elements, prophages, RNAs, and REPINs

**DOI:** 10.1101/2024.07.17.603846

**Authors:** Tue Kjærgaard Nielsen, Lars Hestbjerg Hansen

**Affiliations:** Department of Plant and Environmental Sciences, University of Copenhagen, Copenhagen, Denmark

## Abstract

Microbial genomes are continuously being rearranged by mobile genetic elements (MGEs), leading to genetic configurations that may confer novel phenotypic traits such as antibiotic resistance, degradation of novel compounds, or other metabolic features. Standard genomic sequencing provides a snapshot of a genome in one configuration, but this static image does not give insight into the dynamics of genomic evolution and whether MGEs are actively changing a given genome. We applied single-strain mobilome sequencing to *Escherichia coli* K-12 substrain MG1655 under various stress conditions: UV, SDS, nalidixic acid, tetracycline, cetrimide, and copper. Under these conditions, we quantified the activity of a range of genetic elements, including extrachromosomal circular DNA (eccDNA) from IS elements, RNA genes, the UV-inducible e14 prophage, and intergenic repetitive sites (REP). Of the investigated stressors, copper and SDS are among the largest inducers of eccDNA formation from groups of IS elements, while elevated levels of hypothetical RNA/DNA heteroduplexes of ribosomal and transfer RNAs, and Rhs-nuclease proteins are induced under stress various stressors, especially copper and SDS. This approach holds promise for quantifying the genetic response to environmental stress and implications for genome plasticity. The observed mobilization of IS elements upon copper and other stressors helps to explain co-selection of heavy metals with antibiotic resistance genes and MGEs.

## INTRODUCTION

Microbial genomes are continuously being altered by an onslaught of molecular parasites that often impose no apparent benefits to their hosts. However, under selective conditions, these mobile genetic elements (MGEs) can capture accessory genes that provide e.g. antibiotic resistance or a new catabolic pathway to the microbial host. The mobile genetic parasites include insertion sequence elements (IS), unit- and composite transposons, integron cassettes, integrative conjugative elements (ICEs), integrative mobilizable elements (IMEs), prophages, REPINs (REP doublets forming hairpins)^1^, and others^2^. Many of these groups of MGEs form extrachromosomal circular DNA (eccDNA) intermediates during their lifecycles. Many MGEs affect the expression of adjacent genes, such as through the internal promoters in IS genes^3^. The insertion-deletion-excision and duplication activity of IS elements has been shown to lead to the formation of novel genetic operons in overnight cultures, displaying rapid evolution through MGEs^4^. However, the short-term activity of such genetic elements and general genetic dynamics upon cellular stress remain understudied.

In this study, *E. coli* cultures were grown overnight, allowing them to reach stationary phase, before diluting them in fresh media and immediately treating them with one of several treatments to induce stress response. Stationary phase *E. coli* K-12 are already in a specific stress response termed the stationary phase response, where sigma factor RpoS accumulates and mediates transcription^5^. The regulation of RpoS is complex and several small regulatory RNAs are involved that each respond to a signal^5^.

### Cellular stress and genetic response

Many of the treatments applied in this study (Table 1) are known to lead to oxidative and other types of stress, with the notable exception of tetracycline which has been shown to prevent bacterial SOS response in *E. coli* ^6^ but induces it in other bacteria such as *Vibrio*^7^. Sub-MIC concentrations of copper leads to oxidative stress in bacteria and can co-select for antibiotic resistance^8,9^. Cetrimide is a quaternary ammonium antiseptic that interacts with and damages the cytoplasmic membrane but also interacts with intracellular targets, including DNA^10^. They also lead to the formation of reactive oxidative species and subsequent intracellular stress^11^. UV radiation leads to formation of intracellular ROS and oxidative stress^12^. Both nalidixic acid and UV induces SOS response *in E. coli*, but SOS induction requires *recFOR* for UV and *recBC* for nalidixic acid, likely because UV leads to single-stranded DNA and nalidixic acid leads to double-strand breaks^13^. Sodium dodecyl sulfate (SDS) is a detergent and treatment leads to denaturation of cytoplasmic proteins, among other effects including oxidative stress, as shown in yeast^14^. A distinct SDS shock response has been shown in *E. coli* with induction of proteases to cope with misfolded proteins^15,16^.

**Table 1.** Applied treatments and the minimum inhibitory concentration (MIC).

| Treatment | Dosage | Reference MIC | Supplier |
| --- | --- | --- | --- |
| UV from CL-1000 Longwave UV Crosslinker (120 mJ/cm<sup>2</sup>) | 10 seconds at 250 nm | 30 seconds at 250 nm<sup>32</sup> | UVP (AnalytikJena) |
| SDS | 50 mg/ml | 100 mg/ml<sup>33</sup> | Sigma-Aldrich (Cat. 436143-25G) |
| Nalidixic acid | 3 µg/ml | 10 µg/ml<sup>34</sup> | Sigma-Aldrich (Cat. N8878-5G) |
| Tetracycline | 1.5 µg/ml | 2 µg/ml<sup>35</sup> | Sigma-Aldrich (Cat. T3258-5G) |
| Cetrimide | 37 µg/ml | 37 µg/ml<sup>36</sup> | Sigma-Aldrich (Cat. M7635-100G) |
| Copper | 1.3 µg/ml | 1.3 µg/ml<sup>37</sup> | Sigma-Aldrich (Cat. C8027-500G) |
| No treatment control | No treatment | No treatment | NA |

### Mobilome sequencing for eccDNA detection

Many databases and tools aim to enable prediction of MGEs and other genetic parasites by sequence homology to known examples, however these approaches are limited in their scope. E.g. many prophages are difficult to predict with a high certainty, as phages display an enormous genetic diversity and other groups of MGEs may likewise not be represented in the databases. Recently, an approach to detect activated prophages in bacterial supernatants, termed VIP-Seq, was developed which leverages the fact that bacteriophages are enveloped in protein^17^. However, VIP-Seq does not allow for detection of non-protected circular DNA molecules. The mobilome sequencing approach relies on alkaline lysis plasmid DNA extraction followed by digestion of linear DNA by an exonuclease^18–20^. During DNA extraction and handling, circular chromosomal DNA is likely to break into linear fragments of DNA that are digested by the exonuclease. The degree to which circular DNA molecules are digested by an exonuclease is related to their size, with smaller plasmids being better preserved over megaplasmids and chromosomes^20^. The mobilome approach has been used to study extrachromosomal circular DNA in yeast, metagenomic circular DNA in rat cecum^21^, and mobilome DNA from wastewater^22^. Circularity of DNA molecules from the mobilome approach was shown to be 95% accurate, as validated with inverse-type PCR^21^.

We apply single-strain mobilome sequencing to *E. coli* K-12 substrain MG1655 under various stressors. Under UV stress, the UV-activated prophage e14 is induced significantly. Furthermore, five groups of IS elements, several repetitive extragenic palindromic repeats (REP), Rhs-nuclease protein-encoding genes, rRNA, and tRNAs were found to have higher mobilome coverage upon stress. We hypothesize that RNA/DNA heteroduplexes arise from highly transcribed genes during alkaline lysis plasmid DNA extraction and that other regions of DNA are protected from exonuclease digestion by potential protein interactions. While these heteroduplex molecules are likely artifacts, their increased levels in response to stress treatments provide valuable insight.

Circular intermediate DNA from IS elements forms within 5 minutes after being diluted from stationary phase into fresh medium without other stressors. When some stressors, such as copper and SDS, are added this eccDNA is reduced after 5 minutes. However, the same stressors lead to a significant increase of eccDNA formation 60 minutes after exposure, showing a stress-induced mobilization of IS elements. Previous studies have shown an increased level of plasmid conjugation upon metal stress^23^, including copper which also induced activity of IS element in this study. It is established that heavy metals and other pollutants co-select for the transmission of antibiotic resistance genes in bacteria via MGEs^9,24,25^. This study shows that several stressors, especially copper, induces the formation of circular intermediate DNA molecules from several IS elements within only 60 minutes after exposure. These insights can help to explain how MGEs are activate upon stress response and can lead to mobilization of important DNA, such as antibiotic resistance genes^26,27^ or xenobiotics catabolic genes^20,28–30^.

## MATERIALS AND METHODS

### Experimental setup

A 37°C overnight (ON) LB culture of *E. coli* MG1655 with plasmid pOLA52^31^ was diluted 1:10 by adding 1 ml ON culture into 7 sample tubes with 9 ml clean LB (37°C). Each sample tube with diluted *E. coli* was immediately subjected to the treatments shown in Table 1 and placed back at 37°C. At 5-, 20-, and 60-minutes post-exposure, 3 ml was taken for mobilome DNA sequencing (see below).

### Mobilome DNA extraction and sequencing

Plasmid DNA was extracted and exonuclease treated as described previously^20^. Briefly, 3 ml of culture was subjected to alkaline lysis plasmid DNA extraction, using the Plasmid Midi AX kit (A&A Biotechnology, Gdansk, Poland). Following extraction, DNA was subjected to exonuclease treatment (Plasmid-Safe™ ATP-Dependent DNase; LGC Biosearch Technologies, Hoddesdon, UK) for 1 hour at 37°C and 30 minutes at 70°C to inactivate the enzyme. Following incubation, digested DNA was cleaned using DNA Clean & Concentrator-5 (Zymo Research, Irvine, CA, USA). Sequencing libraries were prepared using the Nextera XT protocol and sequenced on an Illumina MiSeq instrument (Illumina Inc., San Diego, CA, USA) with 2×151 bp paired-end reads.

### Mobilome data analyses

For all software usage, default parameters were applied unless otherwise specified. Paired-end Illumina data were trimmed for adapter sequences with Trim Galore (v.0.6.4)^38^. Each treatment (Table 1) was treated individually. Trimmed paired-end reads were mapped to the fasta genome file containing the MG1655 chromosome and the pOLA52 plasmid sequences, using bwa-mem2^39^ (v. 2.2.1). Samtools^40^ (v. 1.13) was used to retain either “proper pairs” (flags -f 0×2 -F 256 -F 2048) or discordantly mapped reads (flag -F 1294). Discordantly mapped reads are read pairs that either exceed the expected paired-read distance (e.g. reads mapping to either end of a eccDNA molecule) and/or read pairs whose orientation is different from the expected (e.g. paired reads mapping to each end of an eccDNA molecule will have the outward orientation rather than inward). Both types of discordant reads can indicate the start/end position of eccDNA and are used later for defining the terminal positions. Clipped reads were extracted with samclip (https://github.com/tseemann/samclip) with max clip length set to 75 bp (half of read length). These represent aligned reads where up to half of the read length does not match, such as in the ends of eccDNA regions, and were used to predict circularity of ROIs. ROIs displaying a signature (resembling two out of phase sine waves) increase in discordantly mapped and clipped reads in the ends are denominated as eccDNA. Five genes that are not expected to form eccDNA (araB, glnA, Qin, rpoB, ydjU) were included as negative controls for discordant terminal repeats (Supplementary. Fig. 1).

The bam mapping files with “proper pairs” data was split up into individual replicon bam files, i.e. a chromosome file and a pOLA52 file. Genetic regions with high mobilome coverage were identified by first filtering away positions with coverage 20 times higher than the median, as these are certain to be regions of interest (ROIs). Of the remaining positions, only those with 2 times higher than the new, corrected, median coverage was retained, to remove low background positions that are sure not to be ROIs. Finally, only positions above the 97.5% interquartile range of mobilome coverage was kept as potential ROIs per replicon. The start and end positions of identified ROIs were extended to nearby (within 200 bases) positions that had a discordant read coverage higher than three. This was done to increase likelihood of including the terminal positions of eccDNA. All start and end positions were manually verified and curated.

The curated ROI positions were masked from the replicon mapping files and average and standard error replicon coverage were calculated without ROIs, to gauge the background replicon coverage (non-eccDNA). The sequences of ROIs were searched against all replicons (chromosome and pOLA52) to identify any ROIs that are present on both replicons and thus have artificially high mobilome coverage stemming from plasmid background. This happens when e.g. an IS element is sitting on both the chromosome and a plasmid. Since circular plasmids are preserved during plasmid DNA extraction and subsequent exonuclease treatment, chromosomal IS elements will get too high coverage from secondary alignments. ROIs were filtered based on the accumulated mean background coverage from replicons they were identified on.

ROIs passing filters were collated for all treatments and time points and genomic positions were manually curated. All mapping files were manually inspected in CLC Genomics Workbench (v. 22.0.2) and any ROIs that were missed by the automated identification were added to the collated list of ROIs. The coverage of all curated ROIs in 100 bp windows were extracted from all samples, including from those samples where they were not at first identified, for comparisons and statistics.

The 100 bp windowed ROI data was imported in R (v. 4.3.1), where the exonuclease efficiency per sample was calculated as:

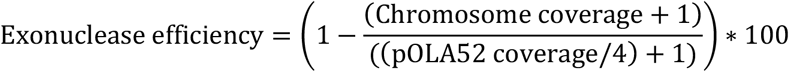

A pseudocount of 1 was added to both replicon (chromosome and pOLA52) median coverages to avoid division by zero, due to zero median chromosome coverage in many samples. The exonuclease efficiency was calculated as the median chromosome coverage (with curated ROIs masked and removed; no ROIs identified on pOLA52) divided by the median pOLA52 coverage (divided by 4 to normalize for assumed 4 pOLA52 copies per cell). This exonuclease efficiency was then used to calculate 100 ROI copies per plasmid in 100 bp windows as:

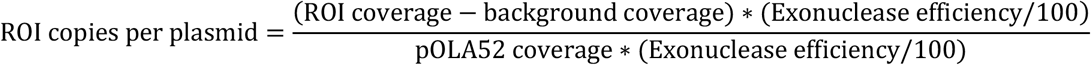

For some 100 bp windows, their coverage was lower than the background coverage of that sample, resulting in negative numbers. These negative numbers were replaced by 0, to reflect that no eccDNA was formed from the given 100 bp ROI windows.

Both ROI adjusted coverage and the exonuclease efficiency-adjusted plasmid coverage could be scaled by the total number of mapped bases, to adjust for differences in sequencing depth between samples, but this will not change the results, as the two adjustments cancel each other out. In a sample with no plasmids for relative abundance comparison, it would be advisable to normalize ROI coverage similar to e.g. RPM (Reads per million mapped reads). However, if a plasmid is present inside the host bacterium, plasmid abundance can be multiplied by exonuclease efficiency, to account for per-sample variation.

Non-parametric Kruskal-Wallis tests, as implemented in the R ‘ggbpubr’ package^41^, on each ROI was used applied to whether ROI data is from the same distribution within each sampling time (5, 20, and 60 minutes). The adjusted relative mobilome coverage increase was compared between the no-treatment control and samples, using post-hoc non-parametric Wilcoxon tests with Bonferroni correction on 100 bp windowed coverage of ROIs.

Miscellaneous features of ROIs were investigated with BLAST at NCBI^42,43^, ISfinder^44^, and RAREFAN^45^. The following R packages were used: ggpubr^41^, ggplot2^46^, and dplyr^47^.

## RESULTS AND DISCUSSION

### Identifying regions of interest

Plasmid DNA extraction and subsequent exonuclease digestion varies in efficiency (Supplementary Fig. 2). Exonuclease efficiency was 98-99.9%, except for cetrimide after 60 minutes (Cet60) that had high chromosomal sequencing coverage. This shows removal of chromosomal DNA and retainment of eccDNA, including pOLA52 for all samples but Cet60. Accordingly, Cet60 was excluded from some analyses. While plasmid copy numbers may vary according to cell growth and environmental stress, the IncX plasmids, such as pOLA52, have a stable low copy number of 3-5 per cell, depending on growth stage^48,49^ and is therefore used as reference for calculating relative abundances of ROIs. After manual curation of computational predictions, 49 ROIs with higher coverage than the expected background remained and were classified into 23 groups (Supplementary Fig. 3). ROIs were merged to groups if they were identical in sequence and were annotated with the same features. E.g. five identical IS*3* regions had elevated mobilome coverage. It may be the case that only one of the five IS*3* form eccDNA, but the reads from that eccDNA maps equally well to all five regions (secondary alignment). The relative abundance of ROIs is normalized to the IncX plasmid pOLA52 that is stably isolated through plasmid DNA extraction and exonuclease treatment. A mean pOLA52 copy number of 4 per cell is assumed (Fig. 1).

**Figure 1.**
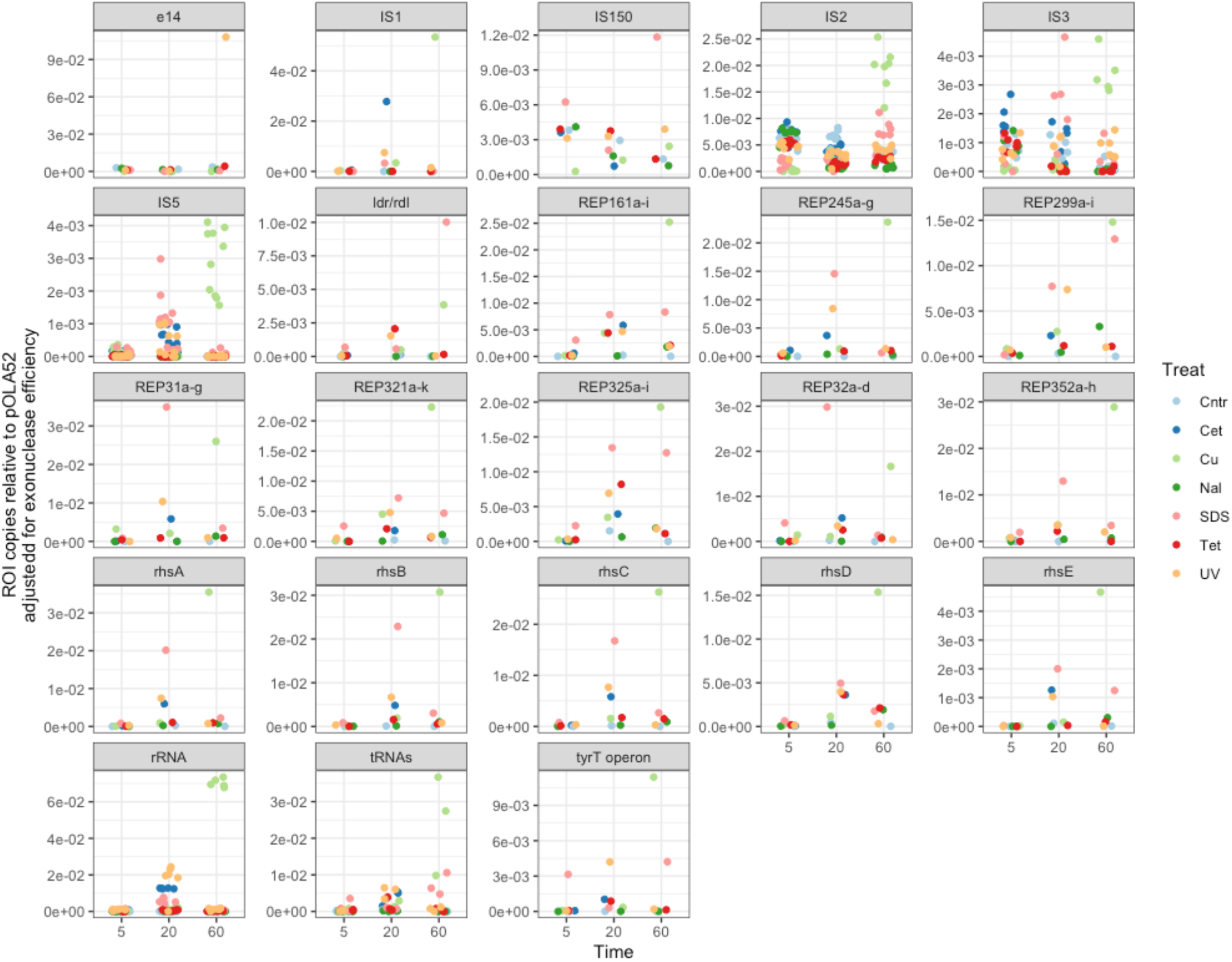
The copies of 23 groups of ROIs normalized by pOLA52 abundance. The background-adjusted coverage of ROIs is divided by the per-sample pOLA52 coverage to obtain per-pOLA52 copy numbers. To obtain the per cell copy numbers of ROIs, an assumed mean pOLA52 copy number of 4 per cell is applied.

### Most stressors decrease coverage of UV-activated prophage e14

As an internal control on the feasibility of the method, we analysed the e14 prophage in response to UV. As expected, the e14 prophage responds to UV exposure^50,51^ after 60 minutes with a 68.92x increase over expected background (Supplementary Fig. 3). However, a 6.22x higher mobilome coverage of e14 is also seen in the control after 60 minutes and 5.67x after 5 minutes and 4.80x higher in nalidixic acid after 5 minutes. This e14 activity is lower in the remaining treatments to a mean 1.03x (SD 0.62), corresponding approximately to no activity over expected background signal. This shows that the activation of e14 is different, depending on the type of stress. It was previously shown that pressure can induce an SOS response in *E. coli* K-12 but not an induction of e14^50^. In the same study, they found that UV induces e14 in approx. 60% of the cells. In our data, the mean normalized e14 abundance in UV after 60 minutes corresponds to 0.11 copies per pOLA52 copy, or 0.027 copies per cell assuming a pOLA52 copy number of 4 (Fig. 1). It is unknown if these copies are equally distributed across cells or concentrated in a subpopulation. This could be resolved in future studies with e.g. single-cell sequencing.

It is likely difficult to replicate the e14 activation levels between studies. The UV lightbulbs of crosslinkers and similar instruments may further vary in dose by age and other parameters. Furthermore, the 60% of cells with e14 activation was measured in late-exponential phase^50^, whereas ours were measured 60 minutes after exposure and in a 1:10 dilution of an overnight culture. As discussed further below, the within-cell genetic dynamics are different according to growth stage and external stimuli.

The no treatment controls display a baseline induction level of the e14 cryptic prophage at 3.1E-03, 1.86E-3, and 3.67E-3 per cell for 5, 20, and 60 minutes, (Fig. 2) showing that e14 has a baseline excision activity, which has been observed previously in a mobilome study^22^.

**Fig. 2.**
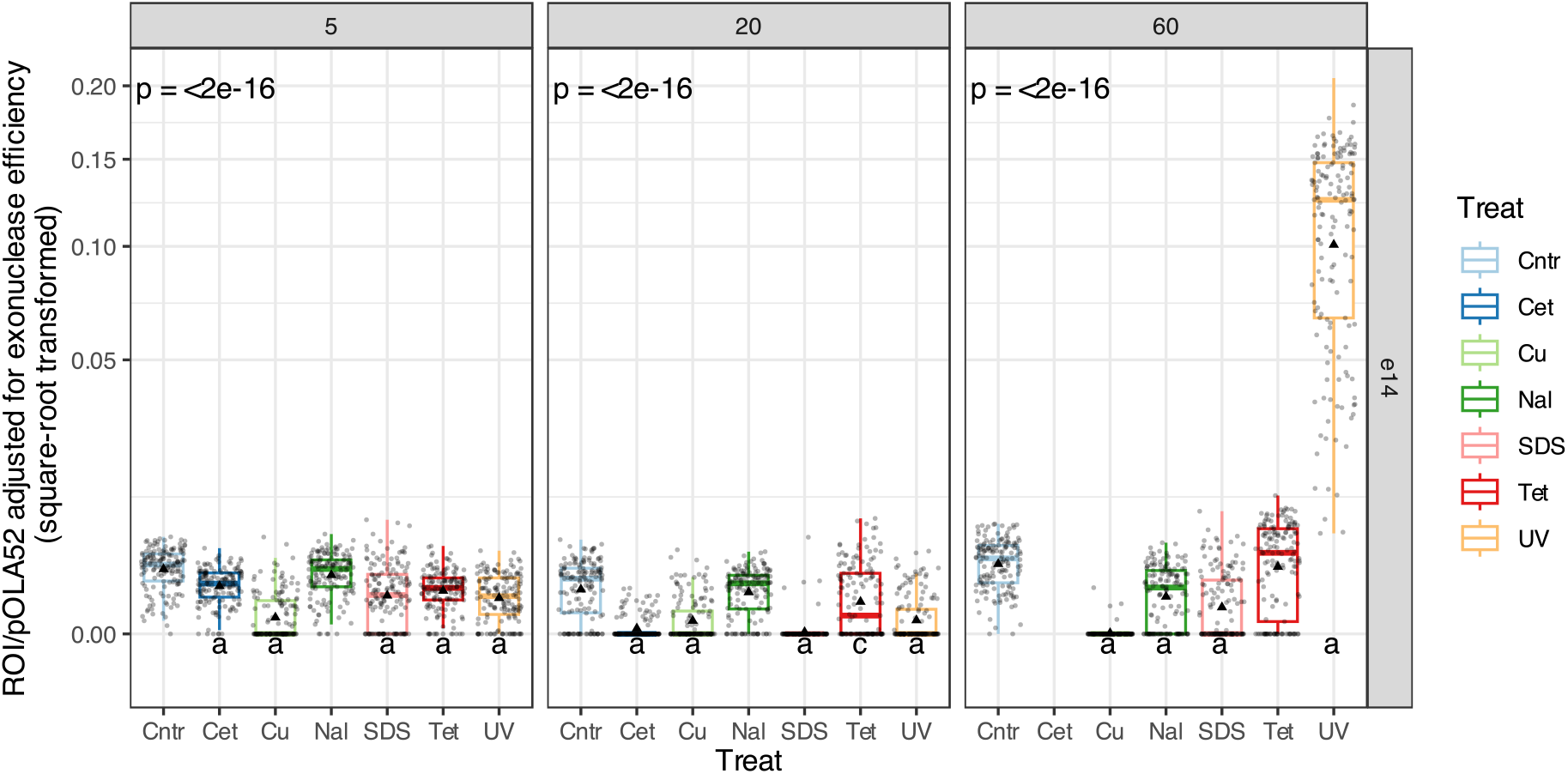
Mobilome coverage of e14 prophage. The p-values in upper left corner are from non-parametric Kruskal-Wallis tests and tests whether data is from the same distribution within each sampling time (5, 20, and 60 minutes). Letters below boxes indicate significance levels from control vs. treatment tested with post-hoc non-parametric Wilcoxon tests with Bonferroni correction on 100 bp windowed coverage; a: p < 0.0001, b: p < 0.001, c: p < 0.01, d: p < 0.05. Boxes show the interquartile range with median as horizontal lines. Whiskers extend to 1.5 times the interquartile range. The mean value is shown as black triangles. Dots show 100 bp window data points. Data has been square-root transformed for plotting.

All treatments except nalidixic acid after 5 and 20 minutes, and tetracycline and UV after 60 minutes decreases the relative abundance of e14 mobilome data (Fig. 2). Mapping of mobilome reads from UV60 shows several spikes of discordant and clipped reads within the e14 chromosomal region (Supplementary Fig. 4). These occur in the ends of the e14 region but also as two higher peaks of unpaired reads internally at even higher depth. This indicates that there might be several eccDNA variants being excised from the chromosome, which was supported by assembly of reads mapping to the e14 region (results not shown). Future studies employing long-read sequencing on eccDNA can unveil e14 eccDNA variants, as individual long-reads, such as those generated by the transposome-based Nanopore rapid sequencing approach. No other prophages on the K-12 chromosome (CP4-6, DLP12, Rac, Qin, CP4-44, PR-X, CPS-53, CPZ-55, CP4-57), were observed to have activity in any of the treatments or time points, although some of these are associated with stress response^52^.

### Stress-induced activation of DDE-type IS elements

Members of the DDE-type IS*3* family of IS elements are known to form circular intermediate molecules (eccDNA) during their replicative transposition^53^. Elevated coverage of discordantly and clipped mapped reads in the terminal regions of the IS*3* family element ROIs, shows that circular DNA is formed from these regions (Supplementary Fig. 5), equivalent to an inverse PCR for eccDNA detection^21^. Clipped ends of reads mapping to terminal regions of IS elements were manually verified in CLC to match the other end of the same IS element, confirming the presence of eccDNA. Terminal inverted repeats were part of the mobilome-sequenced eccDNA region, as analysed with ISfinder^44^. The IS*2*, IS*3*, and IS*150* group IS elements of the DDE-type IS*3* family of transposases are active after 5 minutes of dilution of ON culture with lower relative abundance after 20 and 60 minutes (Supplementary Fig. 3; Figs. 1,3). However, treatment with copper and SDS drops the activity levels of these to the expected background levels, except for IS*150* with SDS at 5 minutes (Supplementary Fig. 3; Fig. 3), indicating that these compounds rapidly decrease the formation of eccDNA molecules of the IS3 family elements. While copper initially represses the eccDNA formation of IS*2*, IS*3*, and IS*150*, it is the strongest inducer of eccDNA formation from IS*2*, IS*3*, IS*5*, and IS2 element after 60 minutes (Fig. 3). Similarly, SDS initially represses some IS elements after 5 minutes, but is inducing eccDNA formation after 60 minutes. Finally, UV induces eccDNA from IS*3*, IS*150*, and IS*1* after 60 minutes. Like e14 ROI, nalidixic acid has no effect on the IS*2*, IS*3*, and IS*150* elements after 5 minutes but is repressing the eccDNA formation of some IS elements after 20 and 60 minutes (Fig. 3), showing that the effect of nalidixic acid is slower than the other treatments. After adjusting for exonuclease efficiency, the IS*1* group is the most abundant with 5.40E-2 and 2.81 copies per pOLA52 for Cu60 and Cet20, respectively, (Fig. 3).

**Figure 3.**
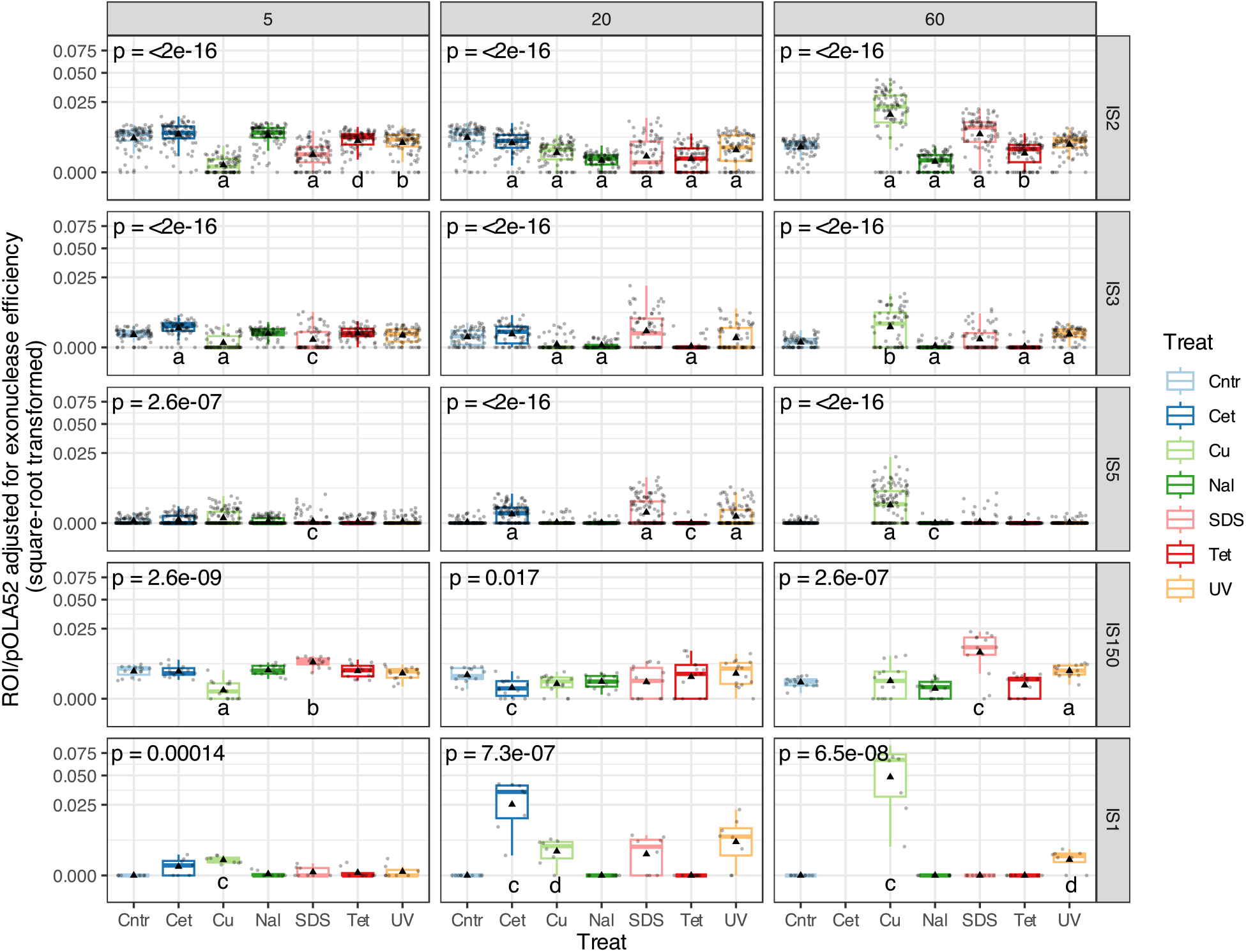
Mobilome coverage of IS element ROIs. The p-values in upper left corner are from non-parametric Kruskal-Wallis tests and tests whether data is from the same distribution within each sampling time (5, 20, and 60 minutes). Letters below boxes indicate significance levels from control vs. treatment tested with post-hoc non-parametric Wilcoxon tests with Bonferroni correction on 100 bp windowed coverage; a: p < 0.0001, b: p < 0.001, c: p < 0.01, d: p < 0.05. Boxes show the interquartile range with median as horizontal lines. Whiskers extend to 1.5 times the interquartile range. The mean value is shown as black triangles. Dots show 100 bp window data points. Data has been square-root transformed for plotting.

The coverage of eccDNA from IS*3* family transposases of the IS*2*, IS*3*, and IS*150* group is generally going down after 5 minutes in no treatment control (Supplementary Fig. 3; Figs. 1,3). It is unknown whether this is due to a lower formation rate of eccDNA, a dilution effect from bacterial cell division, or degradation of eccDNA by bacterial nucleases as a way of reducing the number of intracellular DNA molecules that may lead to detrimental genetic changes. It is unlikely that the lower observation of eccDNA from these MGEs after 20 and 60 minutes is due to successful incorporations into plasmid or chromosome. An experiment over 50,000 generations of *E. coli* showed that IS*150* on average has 9.4 new insertions^54^. For comparison, given a generation time of 20 minutes, no eccDNA from IS elements observed in this study should lead to insertion.

The IS*1* family element is, like the IS*3* family transposase, a DDE-type copy-out-paste-in transposase that has been shown to form circular intermediates (eccDNA) after excision^55,56^. An unexpected mapping signature of discordant and clipped reads for the IS*1F* on the MG1655 (version U00096.3) chromosome, shows that eccDNA of IS*1* elements may be formed from a smaller genetic sequence than annotated in the reference genome (Supplementary Fig. 5). The inverted repeats are furthermore not covered by the increased mobilome coverage or the discordant reads. IS*1* has the highest abundance in Cu60 at 5.40E-2 copies per pOLA52, followed by Cet60 at 2.81E-2 (Fig. 3). IS*1* has a base excision level that is enhanced by the presence of the insertion sequence excision enhancer protein (IEE)^55^ and may furthermore transpose via several alternative pathways that include deletions upon transposition^57^. This is supported by the unresolved eccDNA structure of IS*1* presented in this study, which warrants further attention in future studies.

After adjusting for exonuclease efficiency, the IS*5* family elements are induced by copper after 60 minutes to a per-pOLA52 abundance at approx. 3.26E-3 copies (Fig. 3). While IS*5* elements are DDE type transposases, their transposition mechanism is unknown. Although IS*5* shows a signature discordant read mapping profile, indicating eccDNA formation (Supplementary. Fig. 6), no read pairs overlapping the ends of IS*5* elements could be manually identified. The potential circularity of IS*5* transposition intermediates remain inconclusive. In response to starvation and antibiotic stress, IS*5* has been shown to be activated and inserted in specific sites in the *E. coli* chromosome (*glpFK* operon), enabling improved utilization of glycerol as growth substrate. This stress-induced insertion of IS*5* is associated with a specific DNA structure called superhelical stress-induced duplex destabilization (SIDD)^58^. The IEE stimulates the excision and eccDNA formation of several elements, including the IS*1* and IS*3* family elements, although IS*5* elements were not found to be excised in the presence of IEE^55^. It was proposed that IEE stimulates excision of IS elements that form transposon circles and transpose in a copy-out-paste-in mechanism^55^, although circularity could not be confirmed in this study.

The five groups of IS elements that show elevated mobilome coverage represent only a handful of the diversity of IS elements on the *E. coli* K-12 chromosome. Some, such as IS*30* and IS*As1*, may form even more rare eccDNA molecules than the IS*5* elements here. This is possibly the case for copper after 60 minutes (not shown). Others, like the ISNCY that have been shown to form eccDNA^20^, may not be active under the tested treatments and conditions. Ultimately, other IS elements may not form eccDNA during transposition and are thus not detectable with this method. The fate of the observed IS eccDNA is unknown and it is unlikely that all molecules will be inserted, considering previously observed insertion rates^54^. In the eccDNA from many IS elements, a strong transient promoter is formed from a -35 promoter element in the right end of the IS and a -10 promoter element in the left end. This strong promoter is thought to drive transposase synthesis and commit the eccDNA IS element towards genomic insertion^3^. Future studies should investigate complete genomes after similar exposure and look for genome variants that have new insertions of the numerous eccDNA IS elements observed here.

### Repetitive Extragenic Palindrome (REP) regions respond to treatments

Several intergenic sites were found to be protected from exonuclease digestion, resulting in elevated coverage, especially under SDS and copper stress after 20 and 60 minutes (Fig. 4). These correspond to the Repetitive Extragenic Palindrome (REP) regions REP31a-g, REP32a-d, REP161a-i, REP245a-g, REP299a-i, REP321a-k, REP325a-i, and REP352a-h, as annotated in EcoCyc^59^ for the *E. coli* K-12 version U00096.3. Only REP299a-i is located within an operon (*rhaBAD*), while REP352a-h overlaps with 2 nt with the yjjV gene encoding a putative metal-dependent hydrolase. REP sequences are typically 20 to 40 nt long and are often occurring in tandem inverted repeats separated by linkers in clusters called bacterial interspersed mosaic elements (BIMEs)^60^. There are 355 REP elements in the K-12 chromosome^61^, of which the herein described with elevated mobilome coverage only make up a handful.

**Figure 4.**
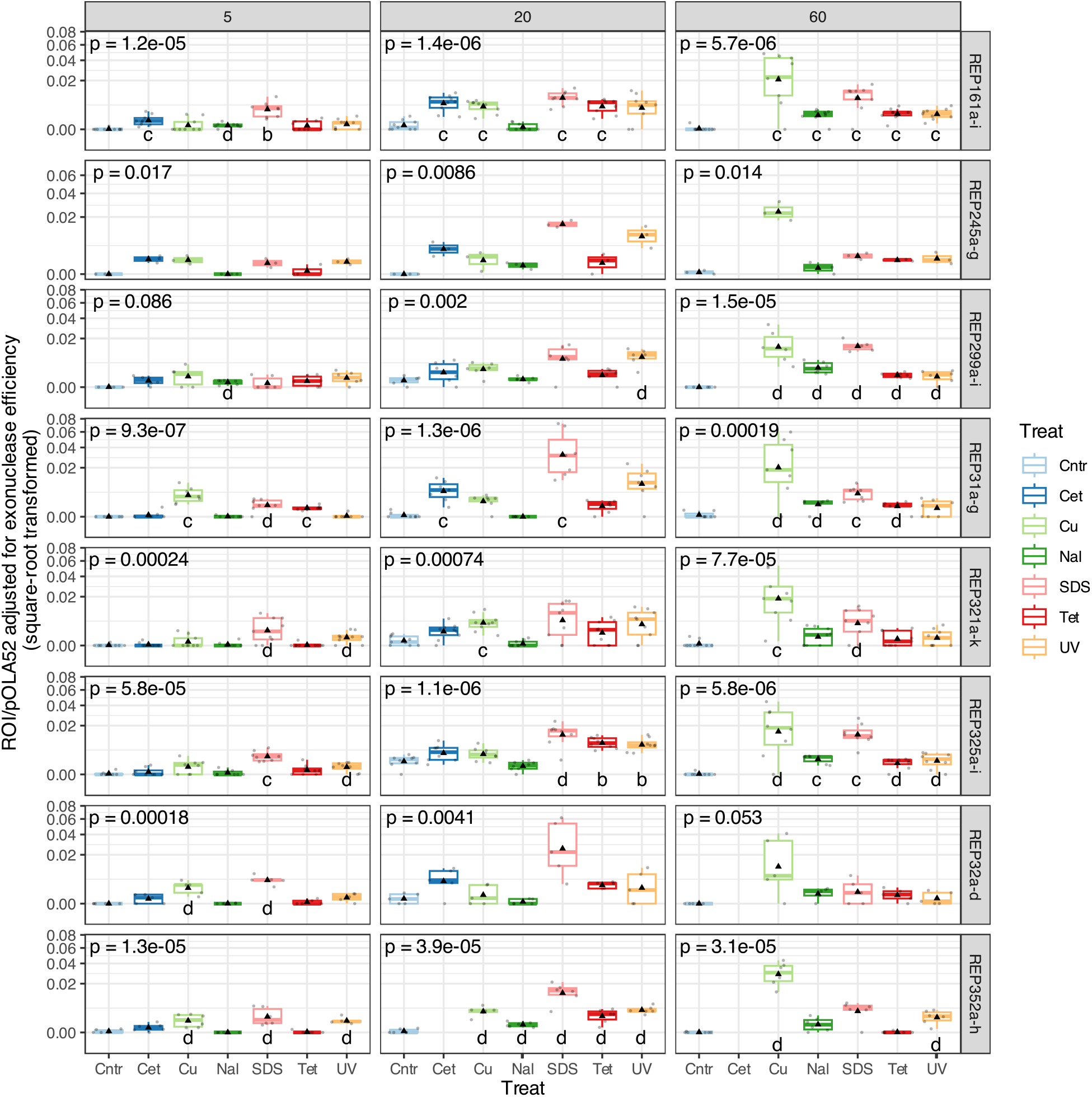
Mobilome coverage of intergenic REP ROIs. The p-values in upper left corner are from non-parametric Kruskal-Wallis tests and tests whether data is from the same distribution within each sampling time (5, 20, and 60 minutes). Letters below boxes indicate significance levels from control vs. treatment tested with post-hoc non-parametric Wilcoxon tests with Bonferroni correction on 100 bp windowed coverage; a: p < 0.0001, b: p < 0.001, c: p < 0.01, d: p < 0.05. Boxes show the interquartile range with median as horizontal lines. Whiskers extend to 1.5 times the interquartile range. The mean value is shown as black triangles. Dots show 100 bp window data points. Data has been square-root transformed for plotting.

REP elements have diverse functions, and some REP sequences are bound by DNA polymerase I, resulting in protection from exonuclease digestion^61^. This has only been shown for REP315 and REP316 in the intergenic regions between *malE/malF* and *lamB/malM* ^61^. Other REP elements are involved in mRNA stability, where they are transcribed together with their upstream genes forming stem-loops in the 3’ UTRs of mRNA, leading to stalling of ribosomes before the stop codon and subsequent degradation of mRNA and translated protein^62^. This gene expression regulation based on ribosome stalling is leaky, with approx. 30% of encoded protein still being completely translated, likely through opening of REP stem-loops on mRNA via RNA helicases^62^. It has been shown that REP-mediated downregulation of translation is reversed by environmental stress conditions, including UV, under which RNA helicases are upregulated. The result is an immediate upregulation of translation of genes that are normally downregulated by REP sequences^62^. This regulatory function may pertain to half of all REP sequences in *E. coli*^62^, since these are located within 15 nt of an upstream gene, which is a hard distance limit for REP-mediated regulation^62^.

All REP sequences in the *E. coli* K-12 chromosome (U00096.3) were downloaded from EcoCyc^59^ and an UPGMA tree (not shown) was made in CLC Genomics Workbench v22.0.2 (Qiagen). All 8 REP regions with elevated coverage contain 4 to 8 imperfect repeats and the repeats are spread out across the entire tree of 697 individual repeats (from 355 REP regions). There is therefore no sequence-based reason why these 8 REP elements have elevated mobilome coverage and not any of the remaining hundreds of REP elements. Of the 8 REP elements, REP299, REP321, REP325, and REP352 are within the 15 nt limit of an upstream gene, required for regulation of protein translation^62^. However, there is no apparent reason why a stress response would lead to increased mobilome coverage of REP elements, based on the model of RNA helicase unwinding of REP stem loops on mRNA for increased protein translation^62^. The transposome from the Nextera XT kit should not attack mRNA – especially not the unwound REP ssRNA stem loops. While 4 out of 8 REP regions with elevated coverage are within the required 15 nt of an upstream gene for translation regulation, we propose that the elevated mobilome coverage of REP elements stems from another mechanism.

### REP exonuclease protection by hypothetical DNA:protein interaction

The domesticated and vertically inherited IS*200*-like REP-associated tyrosine transposases (RAYTs) are, in contrast to most other transposons, typically only found at a single copy per genome, as they do not copy themselves during transposition^1^. RAYTs interact with REPs arranged in pairs forming approx. 100 bp REPINs (REP doublets forming hairpINs) and lead to the multiplication and dispersion of REPs on the chromosome^63^. The function of RAYT is unknown but may include modulate the functions of REPs, as described above. A CRISPR-like behaviour of REPIN-RAYT has also been suggested with the palindromic REPIN structure used as bait for invading DNA that is then inactivated or degraded via endonuclease activity of RAYT^63^. The duplication rate of REPINs is at least 3 orders of magnitude slower than that of IS elements^1^. They duplicate once every 100 million host-cell generations with only 0.2 mutations during this time. In *E. coli* that have lost the RAYT gene, REPIN population numbers have been reduced, indicating that RAYTs are involved in REPIN duplication^1^. Furthermore, the density of REPINs scale linearly with chromosome size, unless the RAYT gene is lost, reflecting an organism responding to niche space. Given their hypothetical endosymbiont-like behaviour, REPINs must confer a fitness benefit to the bacterial host, although it is not clear how. Several functions have been proposed, including mRNA stability, gene expression modulation, localized effects on mutation rates, and genome condensation and structure^1^.

Of the 8 REP regions with elevated mobilome coverage, 7 (REP31a-g, REP32a-d, REP161a-i, REP245a-g, REP299a-i, REP321a-k, and REP325a-i) were predicted as RAYT-associated REPINs by RAREFAN^45^. Here, we show that multiple stressors affect the proportion of REPIN sequence that is protected from exonuclease digestion (Fig. 4). It is possible that this protection stems from the REPINs interacting with RAYT proteins and the increase in coverage is therefore an indirect reflection of RAYT activity. RAYTs are likely derived from the IS*200* tyrosine (Y1)-transposase of the HUH superfamily of ssDNA nucleases that forms eccDNA during transposition. The RAYT gene did not display elevated mobilome coverage in any of the samples, showing that there is no eccDNA from RAYT, supporting previous findings^1^.

The REP325a-i elements encode nucleoid-associated non-coding RNA (naRNA) which is involved in nucleoid DNA condensation through formation of cruciform DNA structures in coordination with the HU protein^64^. Bacterial chromosome topology generally changes in response to environmental stress, including oxidative stress^65^, which is partially mediated by the HU protein binding to DNA to protect it from reactive oxygen species^66^. We propose that our data on REP325a-i, and possibly other intergenic REP regions with elevated mobilome coverage (Fig. 4), reflects chromosomal topology changes in response to stress. It was furthermore found that HU has a higher affinity for RNA/DNA heteroduplexes than just DNA or RNA alone^67^, which may also be a factor in case of the naRNA-encoding REP325a-i, as we suspect these originate from alkaline lysis plasmid DNA extraction.

It is unknown whether the elevated mobilome coverage in REPIN regions stems from expansion of repeats, from formation of structural complexes, or from protection from exonuclease digestion by e.g. protein binding to REP sequences. As explained below, the most likely scenario is protection from digestion by binding of a protein to REP sequences, such as the RAYT. Due to the repetitive nature of the REP regions, a few models for BIME amplification have been suggested, including the formation of circular ssDNA BIME intermediates or rolling replication amplification of a cleaved and excised circular BIME^68^. In our experiment, we used the Nextera XT kit (Illumina Inc.) to build sequencing libraries, which relies on a transposome for attacking dsDNA and adding adapters. This is useful for opening small dsDNA circular molecules, but it should not work on ssDNA. If the elevated coverage in the 8 REP regions represents REPIN duplication in response to stress conditions, then the duplication rate is changed from once every 100 million bacterial generation^1^ to as much as 3.57E-2 extra REPIN copies per pOLA52 copy or one in every 112 cells, assuming a copy number of 4 per cell of the IncX plasmid pOLA52.

We manually investigated the potential formation of circular DNA from the 8 REP elements by looking for read pairs that overlap ends of regions, as circular DNA molecules will have (similar to performing a PCR with outward pointing primers). None of the 8 regions showed signs of circularity, indicating that no circular DNA is formed, as suggested by amplification models^68^. This is supported by discordant read mapping in the termini of REP regions (Supplementary. Figs. 6 and 7). Instead we propose that the 8 REP regions are protected by e.g. a protein that interacts with BIMEs, such as RAYT (TnpA_REP_)^69^. The single copy RAYT transposase does not itself form eccDNA during its own activation^1^.

### Stress response effect on rRNA, tRNA, ncRNA, and Rhs nucleases

The applied “tagmentation” transposomes in the Nextera kit for building Illumina libraries can cut and insert DNA adapters for sequencing on RNA/DNA heteroduplex complexes^70^. Coupled with the DNA-denaturing alkaline lysis method used and the lack of discordant read coverage in these regions (Supplementary Text S2), we find it likely that the observed elevated coverage for RNA genes originates from sequencing of RNA/DNA heteroduplex molecules formed during plasmid DNA extraction.

We observed elevated mobilome coverage of several RNA genes, including rRNA operons, tRNAs, and ncRNA, as well five distinct Rhs nuclease-encoding genes in response to several treatments, with copper and SDS imposing the biggest effect after 60 minutes (Supplementary Text S2; Supplementary Figs. 8-12). These are all believed to act in response to cellular stress, further discussed in Supplementary Text S2. RNA molecules are especially vulnerable to reactive oxygen species^71^ and several treatments in this study are known to cause oxidative stress. The rapid degradation of e.g. tRNAs upon oxidative stress is thought to slow down the formation of toxic misfolded proteins. A subsequent upregulation of tRNAs presumably prevents ribosome jamming and RNA-ribosome dissociation, leading to increased tolerance to oxidative stress^71^. While the degradation of tRNAs in response to oxidative stress is well-documented and degraded tRNAs have been suggested as an oxidative stress sensor^72^, ribosomal RNA levels in response to various stressors is less understood^72^. In transcriptomic studies, rRNA is often removed prior to cDNA conversion to enhance the relative abundance of mRNAs^73^. Most studies on stress response based on cDNA will therefore miss the effects on rRNA and tRNA levels.

A region encoding the long direct repeat toxin-antitoxin *ldr/rdl* module has higher mobilome abundance after 60 minutes in copper and SDS treatment (Supplementary Fig. 8). The *ldrABC* genes encode toxic proteins that rapidly kill the host cell. The *rdlABC* in turn encode unstable antisense RNAs that prevent that translation of the *ldr* mRNA. Ldr protein is associated with growth inhibition, reduced cell viability, nucleoid condensation, and reduced global translation^74^. LdrA was shown to cause inhibition of ATP synthesis as its mode of action^75^. It has been hypothesized that LDR and other toxin-antitoxin systems are involved with arrest of cell growth and activity in response to environmental stress and the formation of persister cells^76^. Our data supports this, and we believe that the signal from the ldr/rdl module is from heteroduplex nucleic acid molecules and not eccDNA (Supplementary Fig. 9). Although a transcriptomic profiling of strain K-12 did not find an upregulation of the *ldr/rdl* module in response to various stressors^77^, the purpose of *rdl* antisense RNAs is to bind to other nucleic acids, possibly allowing the complexes to be sequenced, as observed in this study.

The *tyrT* operon region encoding the *rttR* gene, together with protamine-like protein tpr, and tyrosine tRNA genes *tyrVT* has elevated mobilome coverage in cetrimide, tetracycline, and UV samples after 20 minutes and copper and SDS after 60 minutes (Supplementary Fig. 8). The *rttR* gene encodes rtT RNA which is involved with transcript processing of the adjacent *tyrT* and is likely involved in modulation of the stringent response^78^. In *E. coli, rttR* was previously found to upregulated in response to multiple stressors^79^. While our data supports a role of *rttR* in stress response, the mode of action is still unknown.

### DNA-binding proteins potentially inhibit exonuclease digestion

The RhsA-E proteins have a conserved RhsA nuclease domain (BLASTX). These proteins have been shown to degrade DNA, acting as inhibitors in target cells for intercellular competition, while RhsA-producing bacteria are protected by immunity proteins^80^. After only 5 minutes, *rhsA-D* have significantly higher coverage than the control for all treatment except nalidixic acid. This effect is even more pronounced after 20 and 60 minutes (Supplementary Fig. 12). None of the *rhsA-E* ROIs showed elevated coverage of discordant reads, indicating that eccDNA is not formed from these regions (Supplementary Fig. 12). Instead, we hypothesize that the elevated mobilome coverage stems from either DNA protection by an immunity protein or RNA/DNA heteroduplexes. It is not known if Rhs proteins are involved in stress responses, but it might be the case, based on our data. A transcriptomic profiling of strain K-12 under various stressors, including nutrient- and oxygen limitation, trimethoprim and chloramphenicol treatment, and low pH, showed that none of the Rhs protein-encoding genes responded to stress^77^. Another study found that *rhsBCD* were downregulated in a tetracycline-resistant MG1655 under sub-MIC tetracycline stress^73^. However, *rhsD* was upregulated under butyrolactone stress condition^81^ and in our study, this gene has significantly higher mobilome coverage than the control at 20 and 60 minutes, especially for cetrimide and copper (Supplementary Fig. 11). Furthermore, the *rhsE* gene was upregulated under butanol, butyrolactone, furfural, itaconic acid, and succinic acid stress^81^. In this study, we mostly found *rhsE* had mobilome coverage at or below the expected background coverage, with only small increases in some samples (Supplementary Fig. 11). These mixed results indicate that Rhs proteins may have a role in stress response.

## CONCLUSION

This study applied mobilome sequencing to observe the effects of various stressors on *E. coli* K-12 substrain Mg1655. As expected, the UV-inducible prophage e14 is significantly enriched after 60 minutes post UV application. Copper and SDS are strong inducers of eccDNA from several IS elements. Furthermore, these and other stressors induce an increase in coverage of several other regions, although not forming eccDNA, likely related to stress response. These include REP regions, rRNAs-, tRNA-, and sRNA-encoding genes, and Rhs-nuclease-encoding genes, agreeing with previous studies on stress response at a genetic level.

Mobilome sequencing can be used to quantify not only eccDNA from MGEs, but also to investigate genetic responses to environmental conditions. We envision that the approach can be used to better understand stress responses, as well as the genetic evolution of important phenotypic traits, such as antibiotic resistance and degradation of xenobiotics. The observed formation of circular intermediate molecules from IS elements upon stress supports observed co-selection of MGEs and antibiotic resistance genes upon heavy metal exposure. With the methods of this study, we can now quantify some of these dynamics.

