## Supplemental figures and text for "Single-strain mobilome sequencing quantifies bacterial genetic response to stress, including activity of IS elements, prophages, RNAs, and REPINs"

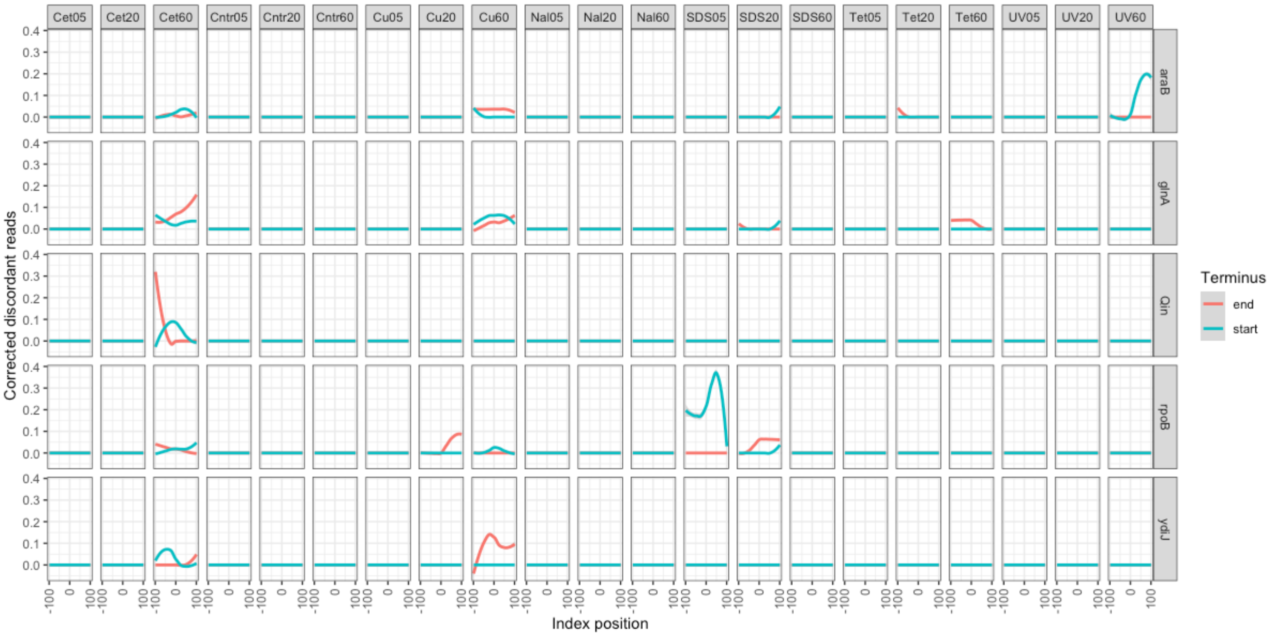


**Supplementary Fig. 1. Discordant reads in the terminal ends of negative control genes not expected to form eccDNA.** 100 bases up- and downstream of the left and right termini of ROIs are included to show that baseline discordant read mapping is absent. The per-ROI discordant reads coverage is normalized against the ROI mean coverage, to compensate for differences in ROI mobilome coverage. A signature discordant read profile resembling two out of phase sine waves shows high discordant reads in both termini of an ROI, indicating the existence of eccDNA. Negative control genes do not display this signature. **See Supp. Fig. 6** for examples of eccDNA signature.

### Supplementary Text S1

#### Exonuclease treatment

Plasmid DNA extraction and subsequent exonuclease digestion can vary in efficiency (**Fig. 1**). A complete exonuclease digestion of chromosomal DNA results in a median chromosome coverage of 0 and an exonuclease efficiency of 100%. Alkaline lysis plasmid extraction and exonuclease treatment failed for sample Cet60, as the low-copy number IncX pOLA52[1] had higher coverage than the chromosome, after adjusting for plasmid copy number (**Fig. 1**). While plasmid copy numbers may vary according to cell growth and environmental stress, the IncX plasmids have a stable low copy number of 3-5 per cell, depending on growth stage[2,3] and is therefore used as reference for calculating relative abundances of ROIs. The relative abundance of ROIs is normalized to the IncX plasmid pOLA52 that is stably isolated through plasmid DNA extraction and exonuclease treatment. A mean pOLA52 copy number of 4 per cell is assumed.

| Treatment | Time | Total mapped bases | Chr. cov. | pOLA52 cov. | Exonuclease eff. |
|---|---|---|---|---|---|
| Cet | 5 | 3.77E+06 | 0 | 2836 | 99.9 |
| Cet | 20 | 2.02E+07 | 3 | 2666 | 99.4 |
| Cet | 60 | 1.82E+08 | 36 | 100 | -42.3 |
| Cntr | 5 | 4.93E+06 | 0 | 6057 | 99.9 |
| Cntr | 20 | 1.63E+07 | 3 | 4462 | 99.6 |
| Cntr | 60 | 8.17E+06 | 1 | 6665 | 99.9 |
| Cu | 5 | 4.59E+06 | 1 | 899 | 99.1 |
| Cu | 20 | 4.05E+07 | 8 | 3122 | 98.8 |
| Cu | 60 | 8.68E+07 | 16 | 1627 | 95.8 |
| Nal | 5 | 6.36E+06 | 0 | 5473 | 99.9 |
| Nal | 20 | 4.09E+07 | 7 | 9269 | 99.7 |
| Nal | 60 | 5.42E+07 | 10 | 6127 | 99.3 |
| SDS | 5 | 1.85E+07 | 4 | 1477 | 98.6 |
| SDS | 20 | 5.53E+07 | 11 | 1246 | 96.2 |
| SDS | 60 | 1.27E+07 | 2 | 707 | 98.3 |
| Tet | 5 | 8.41E+06 | 1 | 3986 | 99.8 |
| Tet | 20 | 6.72E+07 | 14 | 2566 | 97.7 |
| Tet | 60 | 7.30E+07 | 14 | 6044 | 99.0 |
| UV | 5 | 8.32E+06 | 1 | 2688 | 99.7 |
| UV | 20 | 2.58E+07 | 4 | 1240 | 98.4 |
| UV | 60 | 2.31E+07 | 3 | 3885 | 99.6 |

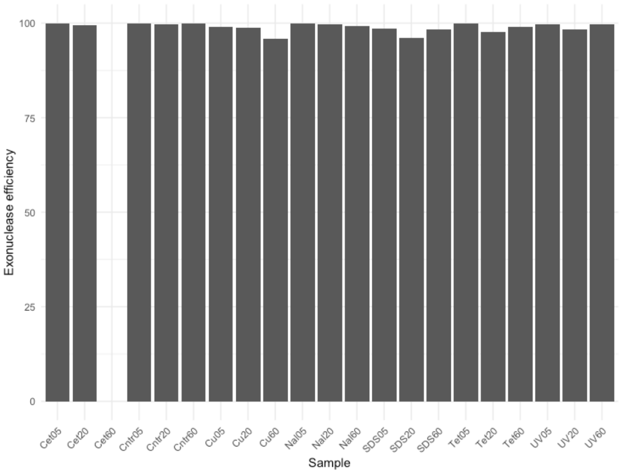


**Supplementary Fig. 2. Replicon coverage and exonuclease efficiency for all samples.** Efficiency is calculated as one minus replicon coverage ratio (chromosome/pOLA52). Both replicons had a pseudocount of 1 added to avoid division by zero. pOLA52 coverage was divided by 4 to account for plasmid copy number.

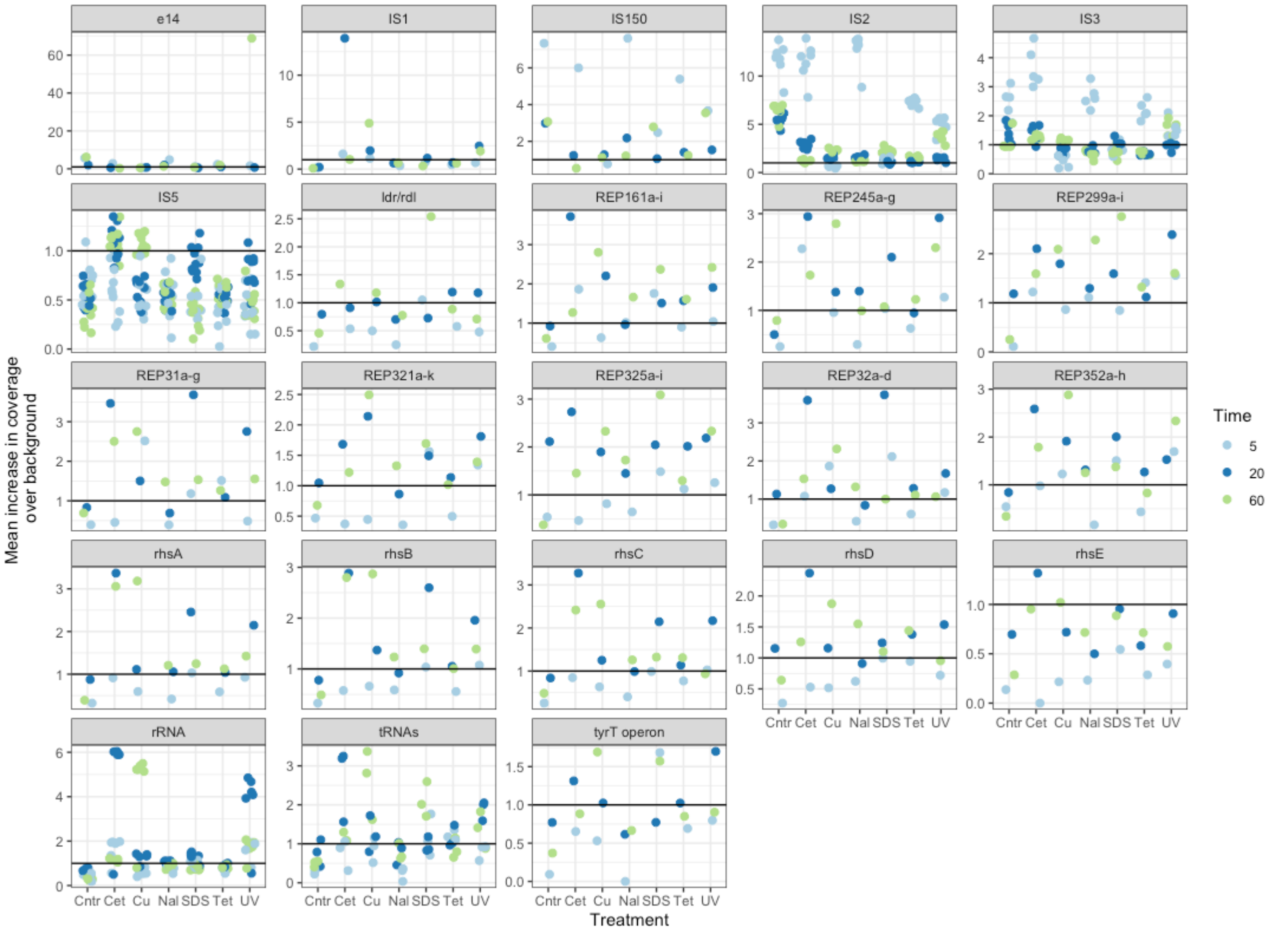


**Supplementary Fig. 3. The increase in coverage of 23 groups of ROIs over the expected background.** These numbers are not normalized by exonuclease efficiency or pOLA52 abundance. A vertical solid line indicates the threshold for over- and underrepresentation of ROIs; i.e. an increase of 1 means that the ROI coverage is the same as the background coverage. The expected coverage is defined as the accumulated mean coverage of replicons with the given ROI plus the standard deviation of the mean. Points within treatments are jittered to avoid overlapping.

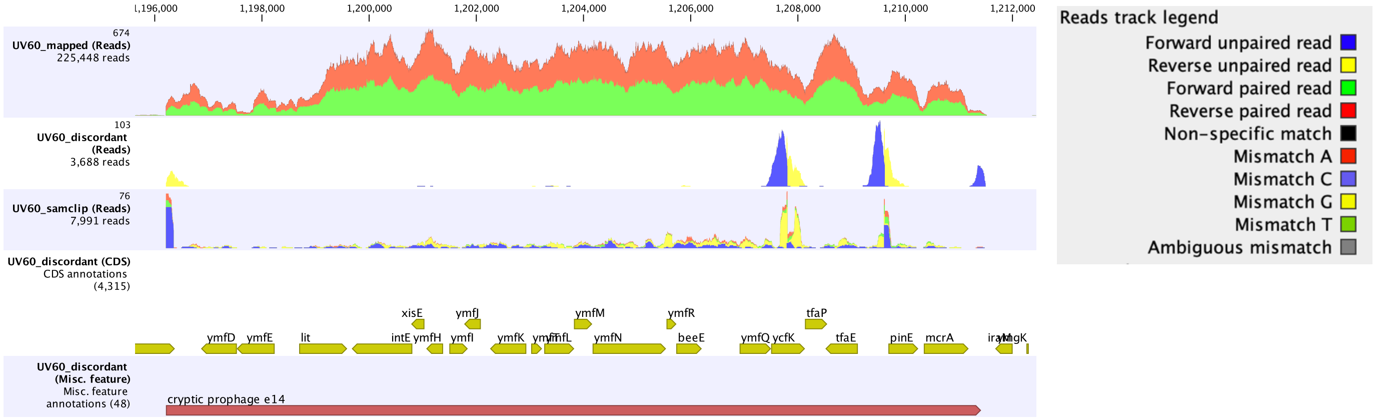


**Supplementary Fig. 4. Read mapping from the e14 prophage region of the UV60 sample.** The first track shows mapping of the total mobilome dataset. The second and third tracks show discordant and clipped reads, indicating several potential terminal ends of the e14 region. The last two tracks show the gene annotations and the e14 annotation from the U00096.3 version of *E. coli* K-12. Legend box on the right shows the type of read alignment. The snapshot is from CLC Genomics Workbench.

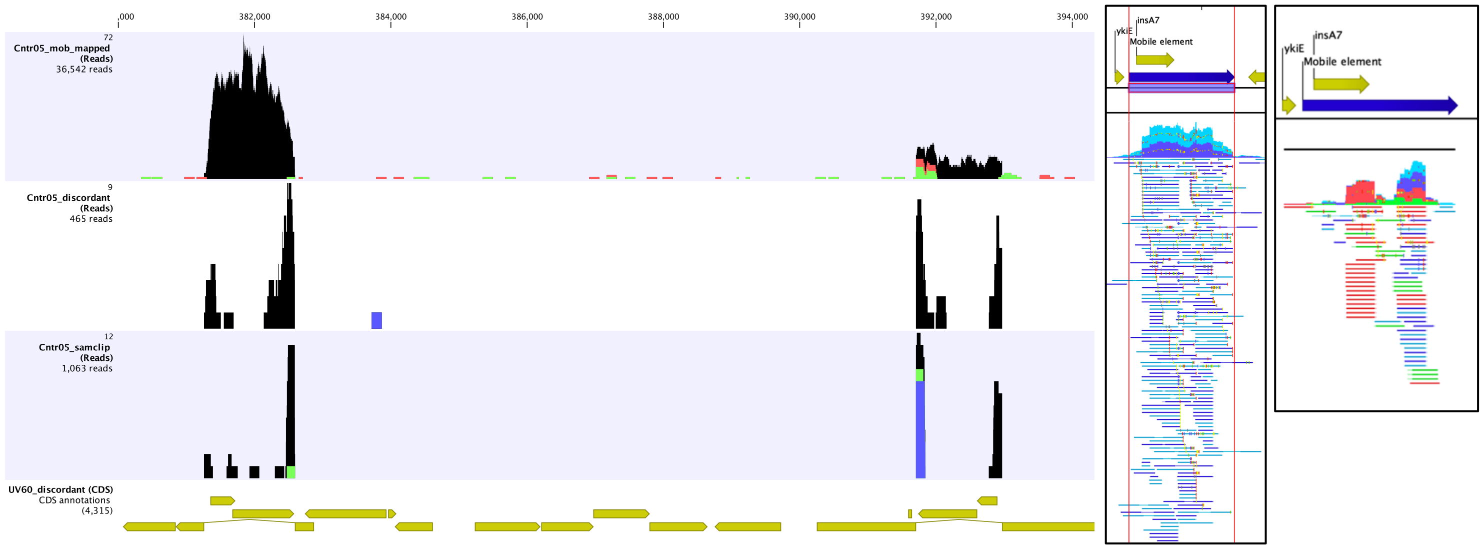


**Supplementary Figure 5. Read mapping from a region with an IS*2* (left) and IS*3* element (right) and zoomed windows (rightmost panels).** The first track shows mapping of the total mobilome dataset. The second and third tracks show discordant and clipped reads, indicating circularity of eccDNA. The last track show the gene annotations from the U00096.3 version of *E. coli* K-12. See Supplementary Fig. 4 for color codes of read alignment. Black alignments indicative non-specific matched reads (secondary alignment), as a consequence of reads mapping equally well to multiple identical IS elements on the K-12 chromosome. The snapshot is from CLC Genomics Workbench.

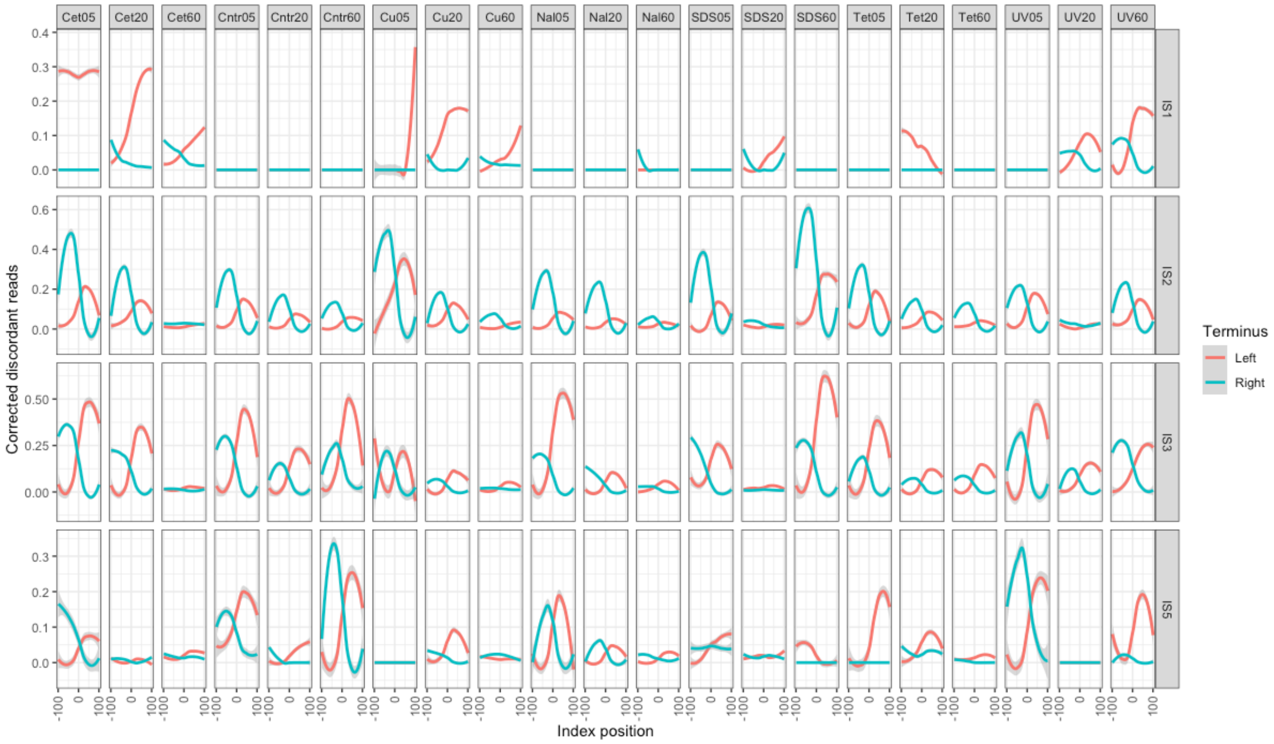


**Supplementary Figure 6. Discordant reads in the terminal ends of IS element ROIs.** 100 bases up- and downstream of the left and right termini of ROIs are included to show that baseline discordant read mapping is absent. The per-ROI discordant reads coverage is normalized against the ROI mean coverage, to compensate for differences in ROI mobilome coverage. A signature discordant read profile resembling two out of phase sine waves shows high discordant reads in both termini of an ROI, indicating the existence of eccDNA.

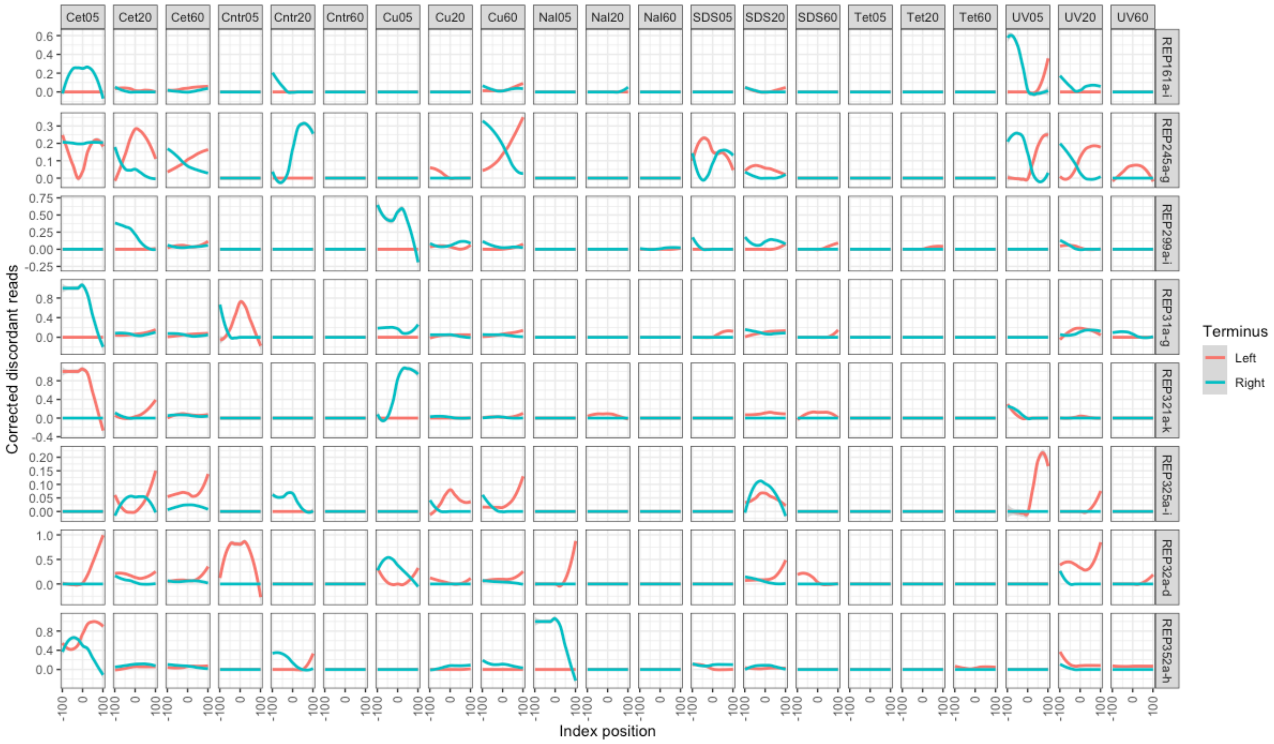


**Supplementary Figure 7. Discordant reads in the terminal ends of intergenic REP ROIs.** 100 bases up- and downstream of the left and right termini of ROIs are included to show that baseline discordant read mapping is absent. The per-ROI discordant reads coverage is normalized against the ROI mean coverage, to compensate for differences in ROI mobilome coverage. A signature discordant read profile resembling two out of phase sine waves shows high discordant reads in both termini of an ROI, indicating the existence of eccDNA.

### Supplementary Text S2

**RNA-encoding regions with elevated mobilome coverage**
We observed elevated mobilome coverage of several RNA genes, including rRNA operons, tRNAs, and ncRNA (**Supplementary Fig. 8**). The applied “tagmentation” transposomes in the Nextera kit for building Illumina libraries can cut and insert DNA adapters for sequencing on RNA/DNA heteroduplex complexes[4]. Coupled with the DNA-denaturing alkaline lysis method used, we find it likely that the observed elevated coverage for RNA genes originates from sequencing of RNA/DNA heteroduplex molecules formed during plasmid DNA extraction. Furthermore, there is no discordant read coverage to support any formation of eccDNA from RNA-encoding regions (**Supplementary Figs. 9 and 10**). However, the biases with which supposed RNA/DNA heteroduplex molecules are formed and sequenced is unknown and quantification is questionable. It can be assumed that the biases are the same across treatments and relative coverage levels should be comparable and, as described below, is likely a function of cellular stress from treatments. RNA molecules are especially vulnerable to reactive oxygen species[5] and several treatments in this study are known to cause oxidative stress. The rapid degradation of e.g. tRNAs upon oxidative stress is thought to slow down the formation of toxic misfolded proteins. A subsequent upregulation of tRNAs presumably prevents ribosome jamming and RNA-ribosome dissociation, leading to increased tolerance to oxidative stress[5].
It was shown that tRNAs are rapidly degraded by cellular enzymes, i.e. RNases[6], upon oxidative stress, with initial decreased tRNA levels 15 and 30 minutes after oxidative stress but increased levels after 90 minutes for some tRNAs, e.g. Ile, Leu, Lys, Met, and Ser[7]. The enhanced tRNA concentrations is associated with maintaining translation and increase resistance to oxidative stress[5,7]. It is thought that only certain types of stress, such as starvation or oxidative stress induces rapid tRNA degradation in E. coli[6]. The data presented here indirectly supports this, as the tRNA levels vary according to treatment and with some ROIs increasing over time (**Supplementary Fig. 8**), assuming data from the tRNA ROIs represents sequencing RNA/DNA heteroduplex.

In our study, we find that some tRNAs have elevated abundance after 20 minutes with cetrimide, copper, SDS, tetracycline, and UV treatment, and furthermore copper, SDS, and UV treatment after 60 minutes (**Supplementary Fig. 8**), although not all tRNAs are elevated, similar to a previous study using tRNA sequencing and qRT-PCR[7]. For Cu60, we find elevated coverage of several tRNAs inside rRNA operons: *ileV*, *alaV*, *gltW*, *alaU-ileU*, *gltU*, *ileT-alaT*, *gltT*, *gltV*, and the following outside rRNA operons: *glnXVWU-metUT-leuW*, *lysTWYZQ-valTZ*, *serX*, *tyrVT*, *valUXY-lysV*, *argQZYV-serV*, *leuVPQ*. While broader than previously reported[7], it is clear that a stress response involving tRNA regulation is complex. It is noteworthy that the same time-dependent regulation of tRNAs is observed in both studies[7] (**Supplementary Fig. 8**).

While the degradation of tRNAs in response to oxidative stress is well-documented and degraded tRNAs have been suggested as an oxidative stress sensor[8], the ribosomal RNA levels in response to various stressors is less understood[8]. In transcriptomic studies, rRNA is often removed prior to cDNA conversion to enhance the relative abundance of mRNAs[9]. Most studies on stress response based on cDNA will therefore miss the effects on rRNA and tRNA levels. During stationary phase and some types of stress, ribosomes go into ribosomal hibernation by the formation of 100S ribosomal dimers. These are rapidly dissociated within one minute of stress relieve, allowing cells to resume growth[10]. The effect of oxidation on the ribosome, depends the affected residues[8] and include impact on translation fidelity, ribosomal stalling, and defective ribosome assembly[5].

In this study, it is unlikely that elevated mobilome coverage of rRNA genes is from sequencing denatured ribosomes, as read mapping of rRNA operons shows uniform coverage of both type-A and type-B rRNA operons[11]. Furthermore, we see no discordant or clipped mapped reads to rRNA operons (**Supplementary Figs. 9-10**), showing that no eccDNA is formed. It is more likely that we are sequencing transcripts from rRNA operons that have formed RNA/DNA heteroduplex molecules with DNA fragments of the operon, as such molecules can be attacked by the Nextera transposome[4]. These would have to be protected from exonuclease digestion. Already 5 minutes after stressors have been applied to cells, there is an increase in rRNA coverage, when compared to the no treatment control (**Supplementary Fig. 8**). While UV is the only treatment that is transient, cells in other treatments will have to adapt quickly to cope. The response in rRNA mobilome coverage is highest for cetrimide and UV treatment after 20 minutes and copper treatment after 60 minutes.

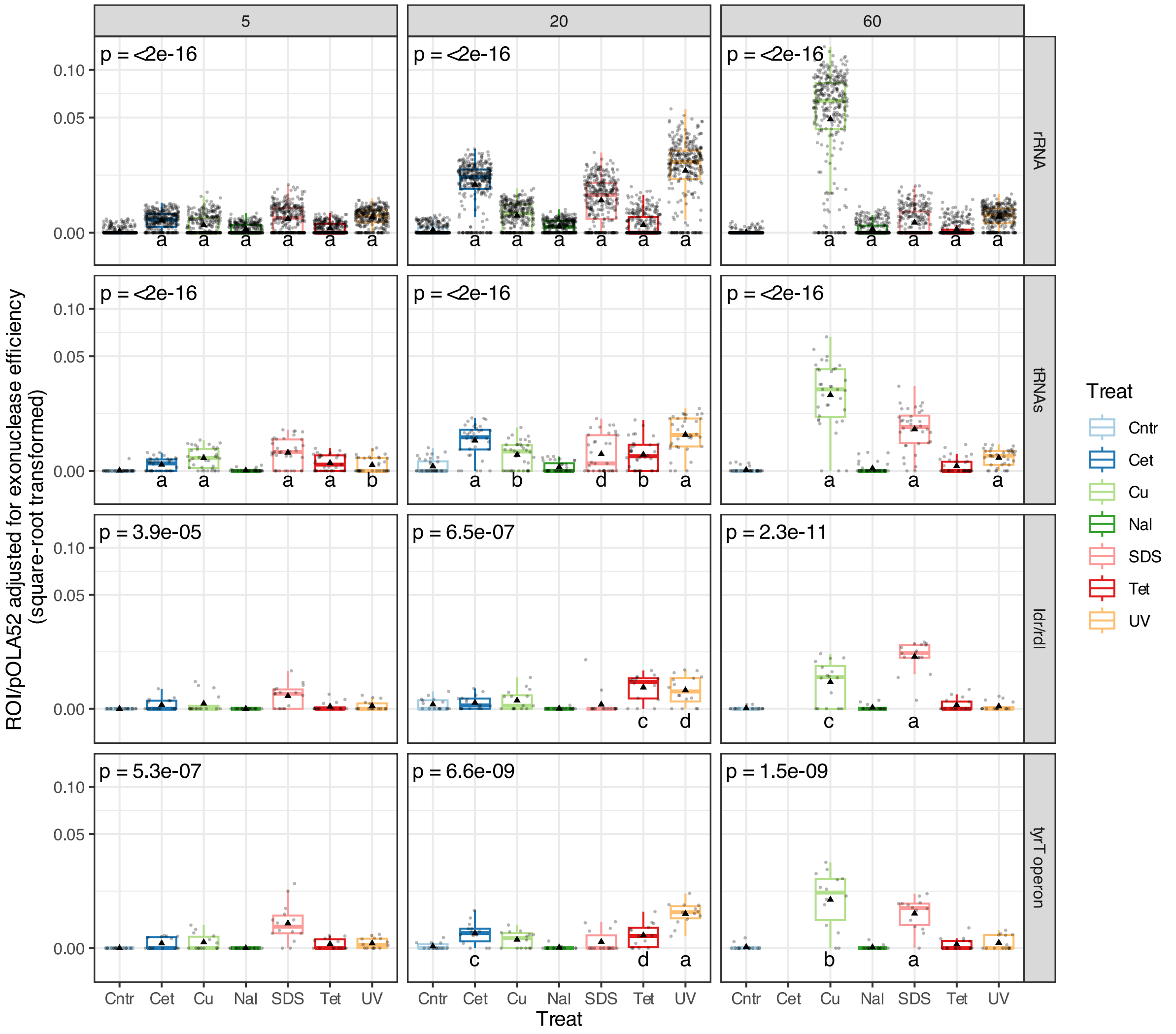


**Supplementary Fig. 8. Mobilome coverage of RNA ROIs**. The p-values in upper left corner are from non-parametric Kruskal-Wallis tests and tests whether data is from the same distribution within each sampling time (5, 20, and 60 minutes). Letters below boxes indicate significance levels from control vs. treatment tested with post-hoc non-parametric Wilcoxon tests with Bonferroni correction on 100 bp windowed coverage; a: $p < 0.0001$, b: $p < 0.001$, c: $p < 0.01$, d: $p < 0.05$. Boxes show the interquartile range with median as horizontal lines. Whiskers extend to 1.5 times the interquartile range. The mean value is shown as black triangles. Dots show 100 bp window data points. Data has been square-root transformed for plotting.

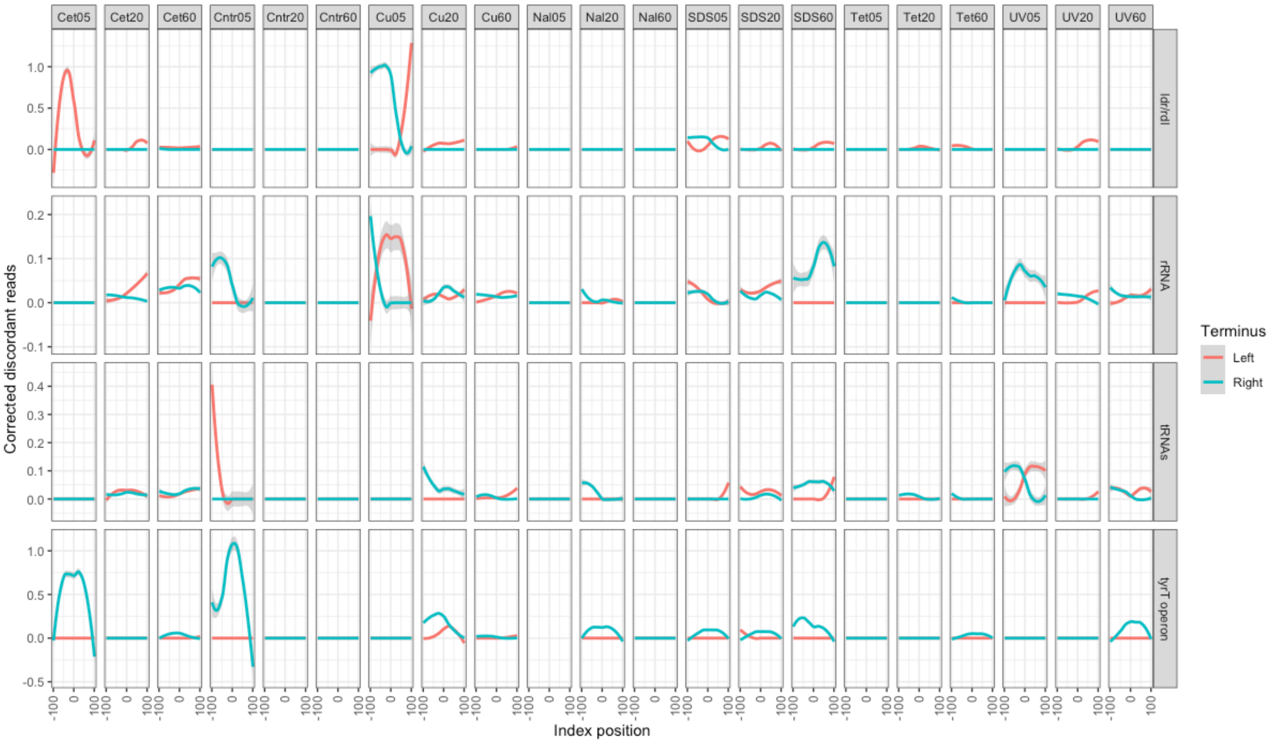


**Supplementary Figure 9. Discordant reads in the terminal ends of RNA-encoding ROIs.** 100 bases up- and downstream of the left and right termini of ROIs are included to show that baseline discordant read mapping is absent. The per-ROI discordant reads coverage is normalized against the ROI mean coverage, to compensate for differences in ROI mobilome coverage. A signature discordant read profile resembling two out of phase sine waves shows high discordant reads in both termini of an ROI, indicating the existence of eccDNA.

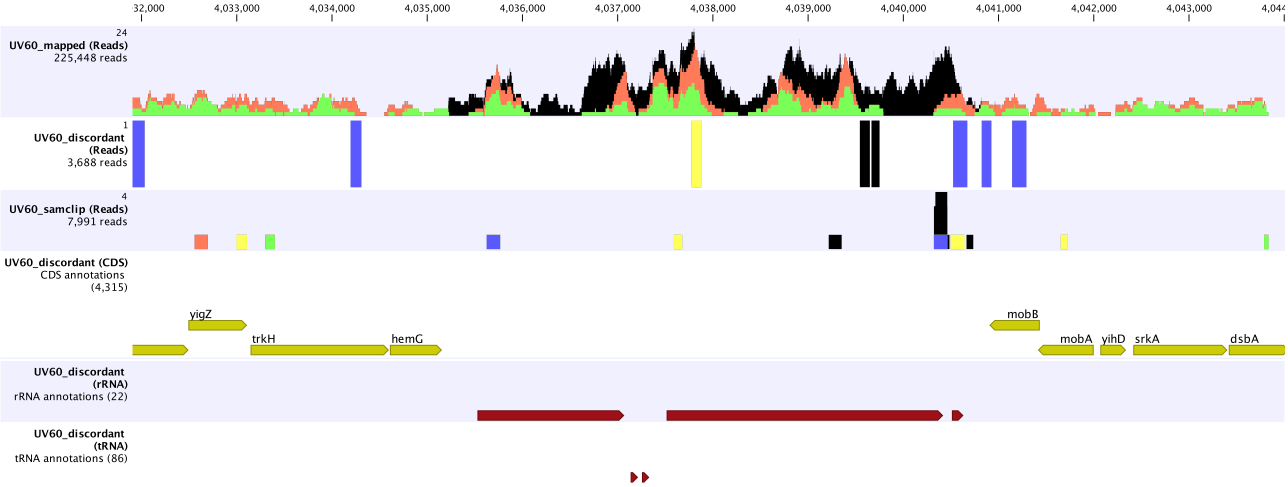


**Supplementary Fig. 10. Read mapping from of an rRNA operon from the UV60 sample.** The first track shows mapping of the total mobilome dataset. The second and third tracks show discordant and clipped reads, indicating circularity of eccDNA. The last tracks show the gene, rRNA, and tRNA annotations from the U00096.3 version of *E. coli* K-12. See Supp. Fig. 4 for color codes of read alignment. Black alignments indicative non-specific matched reads (secondary alignment), as a consequence of reads mapping equally well to multiple similar rRNA operons on the K-12 chromosome. The snapshot is from CLC Genomics Workbench.

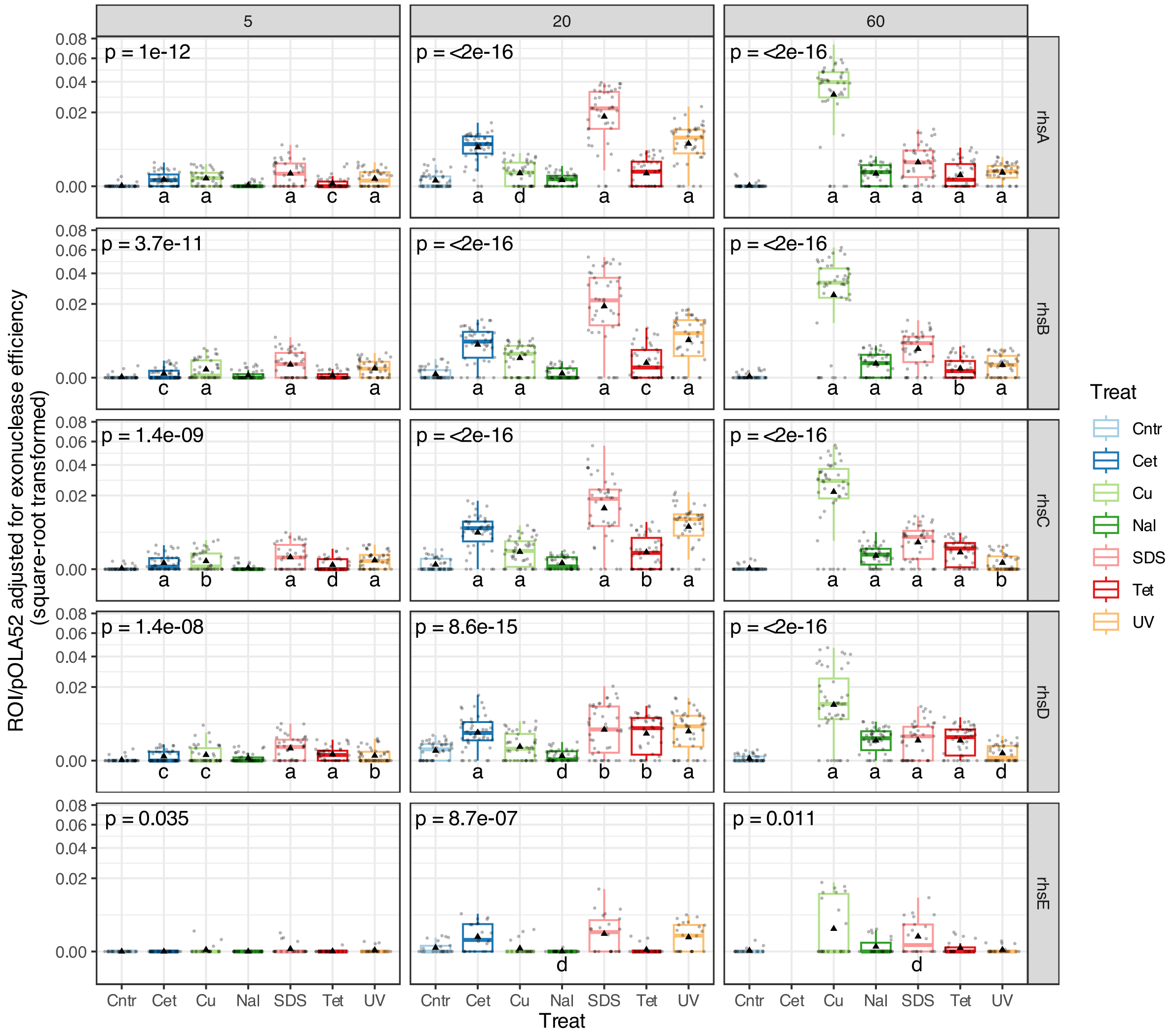


**Supplementary Fig. 11. Mobilome coverage of RHS protein-encoding genes**. The p-values in upper left corner are from non-parametric Kruskal-Wallis tests and tests whether data is from the same distribution within each sampling time (5, 20, and 60 minutes). Letters below boxes indicate significance levels from control vs. treatment tested with post-hoc non-parametric Wilcoxon tests with Bonferroni correction on 100 bp windowed coverage; a: $p < 0.0001$, b: $p < 0.001$, c: $p < 0.01$, d: $p < 0.05$. Boxes show the interquartile range with median as horizontal lines. Whiskers extend to 1.5 times the interquartile range. The mean value is shown as black triangles. Dots show 100 bp window data points. Data has been square-root transformed for plotting.

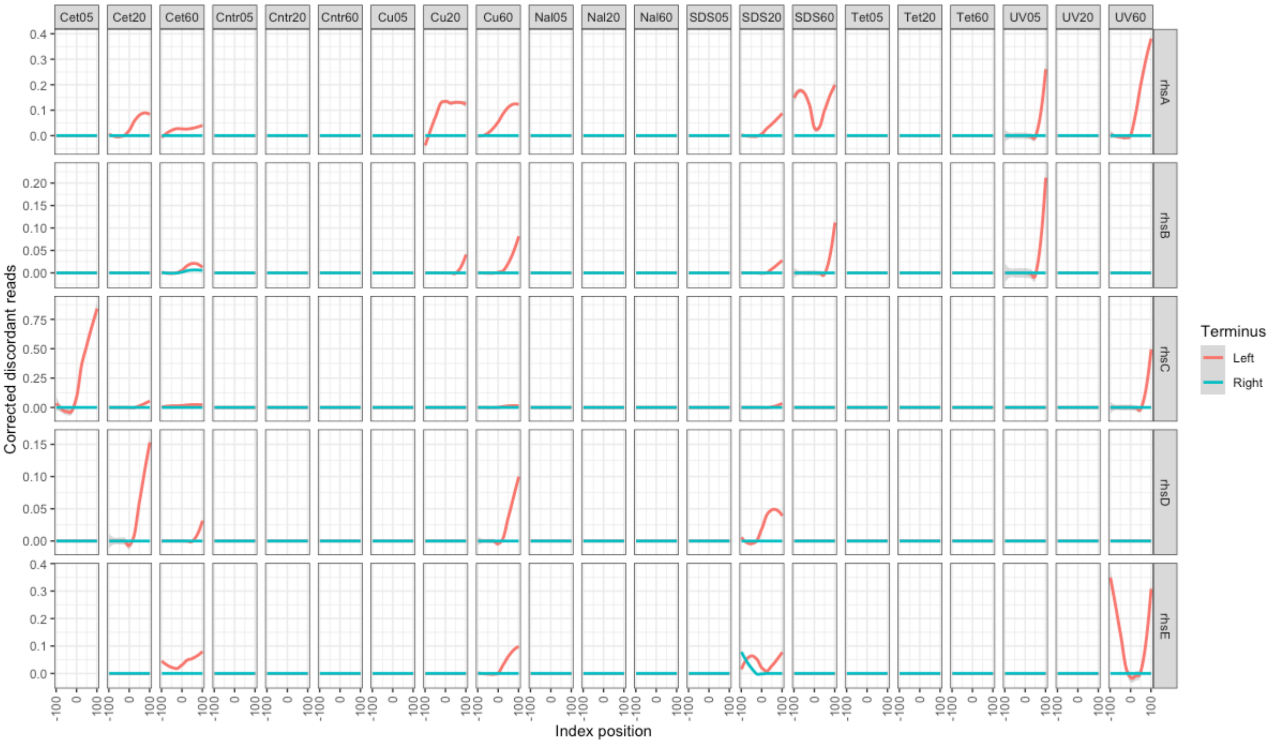


**Supplementary Fig. 12. Discordant reads in the terminal ends of RHS domain-encoding gene ROIs.** 100 bases up- and downstream of the left and right termini of ROIs are included to show that baseline discordant read mapping is absent. The per-ROI discordant reads coverage is normalized against the ROI mean coverage, to compensate for differences in ROI mobilome coverage. A signature discordant read profile resembling two out of phase sine waves shows high discordant reads in both termini of an ROI, indicating the existence of eccDNA.
